# Sex pheromone signal and stability covary with fitness

**DOI:** 10.1101/2021.02.05.429875

**Authors:** Thomas Blankers, Rik Lievers, Camila Plata, Michiel van Wijk, Dennis van Veldhuizen, Astrid T. Groot

## Abstract

If sexual signals are costly to produce or maintain, covariance between signal expression and fitness is expected. This signal-fitness covariance is important evolutionarily, because it can contribute to the maintenance of genetic variation in signal traits, despite selection from mate preferences. Chemical signals, such as moth sex pheromones, have traditionally been assumed to be stereotypical species-recognition signals, but their relationship with fitness is unclear. Here we test the hypothesis that for chemical signals that are primarily used for conspecific mate finding, there is covariation between signal properties and fitness in the noctuid moth *Heliothis subflexa*. Additionally, as moth signals are synthesized *de novo* every night throughout the female’s reproductive life, the maintenance of the signal can be costly. Therefore, we also hypothesized that fitness covaries with signal stability (i.e. the lack of intra-individual variation over time). We measured among- and within-individual variation in pheromone amount and composition as well as fecundity, fertility, and fitness in two independent groups of females that differed in the time in between two consecutive pheromone samples. In both groups, we found reproductive success and longevity to be correlated with pheromone amount, composition, and stability, supporting both our hypotheses. This study is the first to report a correlation between fitness and sex pheromone composition in moths, solidifying previous indications of condition-dependent moth pheromones and highlighting how signal-fitness covariance may contribute to heritable variation in chemical signals both among and within individuals.

## INTRODUCTION

Many sexually reproducing organisms discriminate among potential mates. By selecting a mate, choosing individuals may receive direct benefits, e.g. protection or nutrients, and indirect benefits, e.g. by receiving ‘good’ genes which result in more viable or sexy offspring (1–5). Mate choice commonly occurs through sexual signals, and variation in reproductive success across individuals producing different sexual signals is the basis of sexual selection (3,6,7).

In general, signals under sexual selection are subject to directional selection, as those individuals are chosen that confer the highest direct or indirect benefits. Additionally, in many organisms, sexual signals are used to localize potentially suitable, conspecific mates. These so-called species recognition signals are often under stabilizing selection, because variation in these signals renders them less reliable. Both directional and stabilizing selection are expected to erode genetic variation (8,9). However, sexual signals are shown to have high levels of genetic variance (10,11) and many observations indicate that sexual signals evolve rapidly, diverge early on during speciation, and are important barriers to gene flow among closely related species (6,12–15). To understand how sexual signals evolve, it is important to understand how (genetic) variation in these signals is maintained.

If signals are costly to produce or maintain, their expression and their composition are expected to be correlated with fitness (5,16). Negative correlations between signal and fitness indicate that signal investment trades off with fitness (one-trait trade-offs sensu (17)). Positive correlations between signal expression and fitness are expected when only high-quality senders are able to bear the cost of the signal (two-trait trade-offs sensu (17) and indicate that the signal is condition-dependent (18). Covariation between sexual signal variation and fitness can maintain genetic variation in sexual signals, even in the face of selection (19–22). This is because fitness is the result of the combined effect of many different traits and thus controlled by many different loci (23).

Signal-fitness covariance has been studied mostly in species with acoustic and visual signals, while chemical signals have received much less attention (24,25). This may be because chemical signals, such a sex pheromones, are generally assumed to be biosynthetically cheap (24,26,27) and have traditionally been assumed to be independent of signaler quality (24,28). Specifically, moth sex pheromones have typically been considered as species-recognition signals (29,30) and empirical evidence suggests moth pheromone signal composition is under stabilizing selection (31–35). However, sexual signals probably do not function solely as species recognition or mate choice signals, but rather range along a continuum (20,36,37). This idea is supported by empirical studies that showed a relationship between nutrition and pheromone amount (38), body size and sex pheromone amount (39), and body size and sex pheromone composition (40) in moths, and between nutritional state, age, and parasite load and pheromone composition in beetles (41). However, even though moth sex pheromones have been studied extensively in the past forty to fifty years and the sex pheromone of > 2000 moth species has been identified, very little is known about the relationship between signal variation and fitness or about variation within individuals. Moreover, we lack empirical insight into the relationship between sexual signal variation and fitness for chemical mate attraction in general.

Here, we tested the hypothesis that even for stereotypical species-recognition signals, there is covariation between sexual signal composition and fitness. Since many sexual signals need to be maintained throughout an individual’s reproductive lifetime, and maintenance likely requires continuous investment, we also hypothesized that the ability of an individual to maintain its signal covaries with fitness as well. We tested these hypotheses for the female sex pheromone in the noctuid moth *Heliothis subflexa*. Specifically, we examined how sex pheromone signaling activity (calling activity from hereon), and pheromone amount and composition changed over time.

In moths, older (virgin) females tend to have reduced mating activity and reduced mating success (42–48), and virgin female moths generally keep investing in signaling (49). Thus, prolonged virginity is expected to trigger physiological responses that modulate resource allocation between somatic maintenance and reproduction. We assessed these trade-offs by comparing a biologically realistic delay to first mating (3 days) with an extreme case of prolonging female virginity (8 days), here referred to as ‘early’ and ‘late’ maters, respectively. We then asked whether a) the composition and amount of the pheromone signal in early and later maters covaried with fitness, and b) the stability (within-individual variation) of the pheromone during prolonged virginity was correlated with fitness. We addressed these questions separately in the early and in the late maters to explore the robustness of our findings to a) variation in age and b) the time span over which intra-individual variation was measured. We expected that high-fitness females produced more pheromone compared to low-fitness females, and that maintaining high pheromone amounts and stable pheromone composition trades off with fitness.

## METHODS

### Insects

The laboratory population of *H. subflexa* originated from North Carolina State University and has been reared at the University of Amsterdam since 2011, with occasional exchange between NCSU, Amsterdam, and the Max Planck Institute for Chemical Ecology in Jena to maintain genetic diversity. The rearing was kept at 25°C and 60% relative humidity with 14h:10h light-dark cycle. Larvae were reared on a wheat germ/soy flour-based diet (BioServ Inc., Newark, DE, USA) in 37-mL individual cups. Pupae were separated by sex and checked for emergence every 60 minutes from 2 hours before until 7 hours after the onset of scotophase. Only females that emerged within this timeframe were included in the experiments. After emergence, females were transferred to clear 475-mL observation cups covered with fine mesh gauze. Males were kept in the 37-mL individual cups. Adults were provided with regularly replaced cotton soaked in 10% sugar water.

### Phenotyping

The pheromone of each female was sampled at two time points. The first sample was taken in the first night after emergence. The second sample was taken in the third or eight night after emergence for the ‘early’ and ‘late’ treatment, respectively. Early maters were females that were kept virgin for three days and mated on the fourth day, while late maters were females that were kept virgin for eight days and mated on the ninth day. To avoid possible variation due to differences between sampling time during the night (50–52), care was taken to take the first and second pheromone sample at the same time during the night (no more than 10 minutes difference) and the sex pheromone was always sampled within a narrow time interval during peak calling times (3^rd^ – 6^th^ hour of scotophase (50).

Pheromone was collected from the gland surface from females using optical fibers coated with a 100-μm PDMS (Polymicro Technologies Inc., Phoenix, AZ, USA), as described in detail in (53), where the authors showed strong correspondence between pheromone measurements following the non-invasive method used here and following traditional (lethal) gland extractions. Pheromone glands were extruded and fixed by gently squeezing the abdomen and gently rubbed over a period of 2 minutes. Fibers were then submerged for 1-2 hours and rinsed in 50 μL hexane with 200 ng pentadecane as internal standard and discarded. Pheromone extracts were analyzed by injecting the concentrated samples into a splitless inlet of a 7890A GC (Agilent Technologies, Santa Clara, CA, USA) and integrating the areas under the pheromone peaks, using Agilent ChemStation (version B.04.03).

Throughout this study, we focused on the sex pheromone components that have been shown to be important for male attraction (Z11-16:Ald, Z11-16:OAc, Z11-16:OH, and [Z7-16:Ald + Z9-16:Ald],), as well as the total amount of pheromone. Z7-16:Ald and Z9-16:Ald were difficult to separate by GC and were therefore integrated as one peak (referred to a Z7/Z9-16:Ald), as has been done in previous studies (54). Absolute amounts (in ng) and percentages (of the total amount) of each compound were calculated relative to a 200 ng pentadecane internal standard. Samples containing < 10 ng could not be reliably integrated and were excluded from subsequent analyses. We globally standardized the absolute amount of each of the four pheromone components by subtracting the global average amount across all females and then dividing the mean-centered value by the variance. We then performed principle component analysis on the standardized pheromone measurements, retaining four principle components (PCs). Differences in PC scores between the first and second pheromone samples were tested using a paired student’s t-test, using the statistical package R (55).

### Relationship between fitness and female calling activity

To determine whether calling activity of virgin females increased or decreased over their lifetime or whether it peaked at some point in their reproductive live, we randomly assigned 40 and 51 females to the ‘early mating’ and ‘late mating’ treatments, respectively, and observed the calling activity of each female every night from eclosion until her death. For each female, we measured pupal mass, calling behavior, fecundity, fertility, and lifespan. Pupae with a visible wing pattern were weighed before the start of scotophase (the 10-hour dark phase during the day-night cycle). Extremely small pupae (< 0.1 gram) were discarded. The time spent calling before and after mating was recorded daily every 30 minutes between 1-8 hours after the onset of scotophase. To measure fecundity and fertility, females were mated in 475-mL observation cups with virgin 2-3-day-old males one day after the second pheromone sample was taken. For all mated females, we scored the amount of eggs, the number of hatching larvae, and the life span. To stimulate oviposition, freshly cut *Physalis peruviana* berries were supplied daily onto the gauze. Before the onset of a scotophase, eggs laid in the previous scotophase were collected by transferring the female to a new observation cup. Eggs were counted before emergence, and artificial diet was provided for emerging larvae. Emerged larvae were counted daily. After female death, lifespan was recorded, and mating success was confirmed by checking for the presence of a spermatophore. The percentage of calling females, the per-individual onset of calling and the per-individual duration of calling were visually examined before and after mating. We tested whether early maters differed from late maters in fecundity (overall and per day), fertility (overall and per day), and life span using two-sample t-tests.

### Relationship between fitness and *inter*-individual variation

To test whether fecundity, fertility, and life span variation could be predicted by variation in the pheromone, we fitted generalized linear models with quasi-Poisson error and used a stepwise model selection approach to identify significant predictors and interactions. In the base model, the response was either fecundity, fertility, or life span and the predictors were each of the four PCs as well as the pupal mass of the female, the pupal mass of her mate, and the time spent calling by the female before mating. Only the PCs from pheromone measurements taken on timepoint 2 were used, as these were the samples taken immediately before the matings were set up that were used to measure the individual’s fitness. Model selection was done by first adding interactions among the independent variables and then purging independent variables that did not significantly explain variation in the response. Adding and removing variables and interactions was done using Analysis of Deviance, where significance of an increase/decrease in residual Deviance after removing/adding a variable or interaction was assessed using a Chi-square test. Improvement of model fit was considered significant for p < 0.05. For the final model, the pseudo-R^2^ was calculated as the ratio between the null deviance and 1 minus the residual deviance, similar to calculating the coefficient of determination in linear regression. The pseudo-R^2^ gives the proportion of deviance explained, which is informative about model fit and about the explanatory value of the independent variables (56).

### Relationship between fitness and *intra*-individual variation

To test whether fecundity, fertility, and life span variation could be predicted by the degree to which females kept their pheromone stable, we calculated intra-individual variation in pheromone by taking the absolute difference between PC scores for pheromone measurements taken after 24 hours and PC scores for pheromone measurements taken three or eight days post-eclosion, for early and late maters respectively. We fitted the same generalized linear models (with the same covariates) and employed the same model selection procedure as described above, except that the predictor variables were now the absolute change in PC scores between timepoint 1 and timepoint 2.

### Heritability of stability

To determine the heritability of pheromone composition (PC scores) and stability (difference between PC scores at timepoint one and two), we selected 20 females that covered the range of intra-individual variation observed across all females. Subsequently, we sampled 5-20 daughters per female (N = 240) at both timepoints, i.e. three or eight days post-eclosion for early and late maters, respectively. We used Bayesian Monte Carlo Markov Chain models combined with pedigree information of the two generations of females and their mates. We ran independent models for each principle component (inter-individual variation) or for the change in PC scores between timepoint 1 or 2 (intra-individual variation). The models were implemented with the R package “MCMCglmm” (57) using inverse gamma priors. All MCMC models were run for 1,000,000 iterations, the initial 100,000 samples were discarded (burn-in period) and chains were sampled every 100 iterations (thinning) to reduce autocorrelation. The narrow-sense heritability (*h*^2^) was estimated as the ratio of additive genetic variance (‘animal’ effect) over the sum of all variance compounds (‘animal’ plus ‘unit’ effects).

## RESULTS

### Relationship between fitness and female calling activity

When observing female calling activity, we found that calling effort of all females and duration of individual calling activity initially increased and then decreased after the second or third night (Fig 1A,B). Upon mating, calling activity was almost entirely suspended, after which it slowly recovered. There was no apparent relationship between the time at which females started calling and female age or mating status (Fig S1).

**Fig 1.**
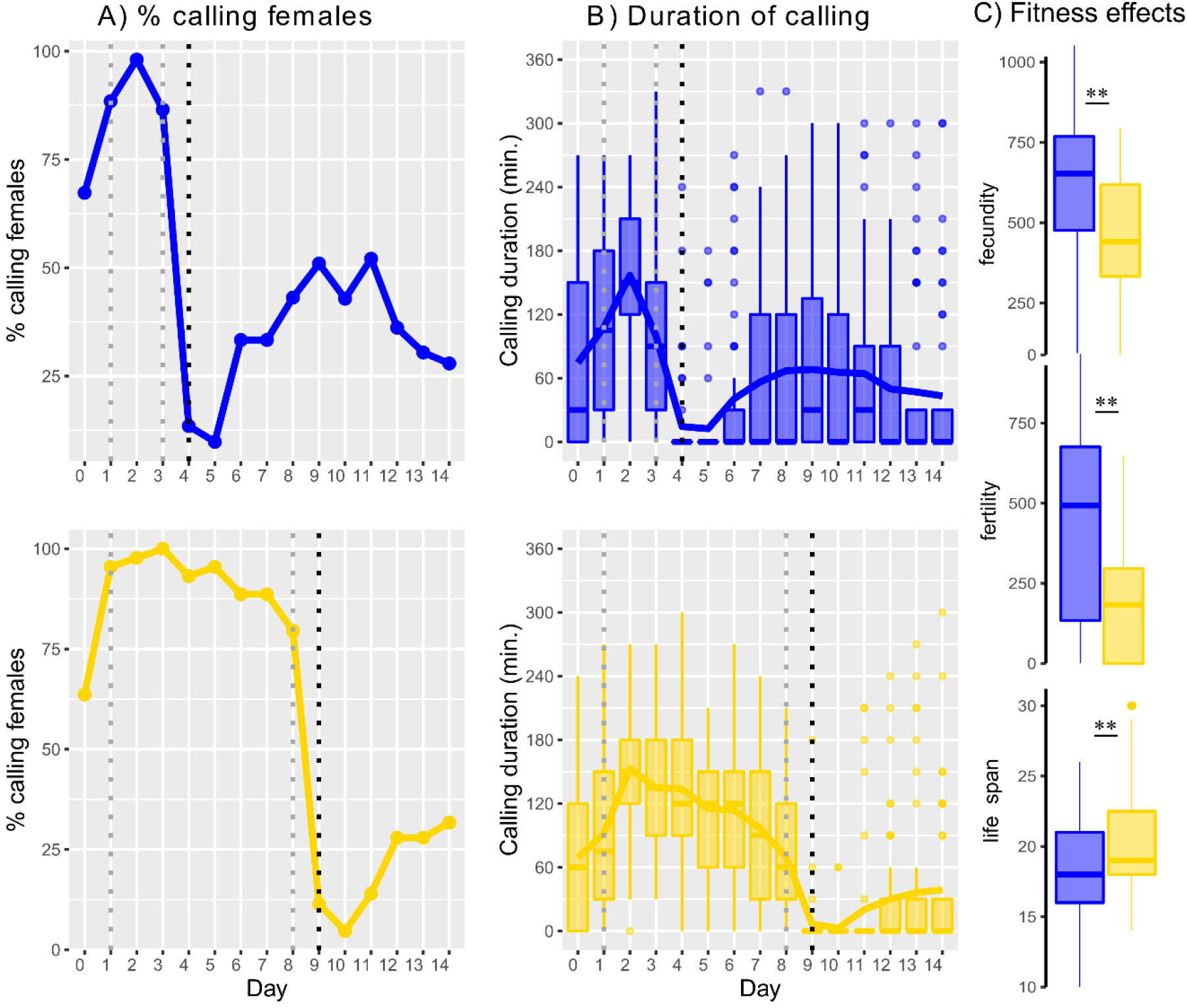
Calling activity and fitness. For both groups, early (blue) and late (yellow) maters, the collective calling effort per night and the distribution of calling duration across females is shown for each night. A) Percent of females calling on any given night, B) Calling duration. Boxes and whiskers show the interquartile ranges and the median, and lines connect mean values for each night. Individual dots show observations > 1.5 times the interquartile range. The dashed, vertical grey and black lines show the days at which pheromone samples were collected and females were mated, respectively. C) Fitness variation within and between early and late maters. **: P < 0.01.

Late maters invested in pheromone calling for an additional five days relative to early maters (Fig 1). Both fecundity and fertility were significantly lower in late maters relative to early maters (Fig 1C). Reduced fecundity was likely due to the ‘lost’ 5 days rather than due to allocation of time and energy, because fecundity per day was not significantly different between early and late maters (46.50 versus 40.96 eggs per day; t = 1.39; P = 0.1669). Fertility per day was lower in late versus early maters (19.00 versus 32.84 larvae per day; t = 3.102; P = 0.0026), indicating that females that mated later in life laid eggs with lower hatching success compared to early maters. We further observed that late-mated females lived on average two days longer than early-mated females (Fig 1C).

### Variation in sex pheromone signal strength and composition

In quantifying the sex pheromone variation using principle component scores for the standardized absolute amounts of four components that are important for male response in *H. subflexa*, we found that the first PC accounted for 60.5% of the total variation among individuals. Since this axis was almost perfectly correlated with the total amount of pheromone (Pearson’s *r* = −0.99; *P* < 0.0001), PC1 thus described variation in the total amount of pheromone. The remaining three PCs described different dimensions of pheromone composition space: higher scores on PC2 (19.8% var. expl.) indicated decreasing amounts of Z11-16:OAc relative to all other compounds, higher scores on PC3 (13.2% var. expl.) indicated increasing amounts of Z11-16:OH relative to Z7/Z9-16:Ald, and higher scores on PC4 (6.54% var. expl.) indicated increasing amounts of the major component, Z11-16:Ald, relative to all other compounds (Table S1; Fig S2). Neither of these three ‘composition’ PCs were correlated with the total amount of pheromone (Pearson’s *r* ranged from −0.05 to 0.08; *P* ranged from 0.3337 to 0.4987) and a significant proportion of the variation in all PC scores was additive genetic variance, except PC1 in the late maters (posterior mode of *h^2^* ranged from 0.17 to 0.58; for PC1 in late maters *h^2^* < 0.01; Fig S3). There was thus no need to calculate log-contrasts for the components to break the unit-sum constraints that trouble the use of relative amounts (58), thereby avoiding inadvertent effects resulting from the choice of the divisor.

With the exception of PC4, all PCs were significantly different between timepoint 1 and timepoint 2, both for early and late maters (Fig 2). This shows that there is intra-individual variation in both amount and composition of the pheromone signal. Over time, pheromone amount decreased, while relative amounts of Z11:16:OAc and Z11-16:OH increased (Fig 2). However, there was substantial variation around the mean direction and magnitude of change, with some individuals showing much more or much less change in the amount and composition of the pheromone over time (Fig 2). This shows that there is variation in the magnitude of intra-individual variation. Estimates of the narrow-sense heritability of intra-individual variation in the PCs were between 0.1 and 0.2, indicating that a significant proportion of the variation is additive genetic variation (Fig S3).

**Fig 2.**
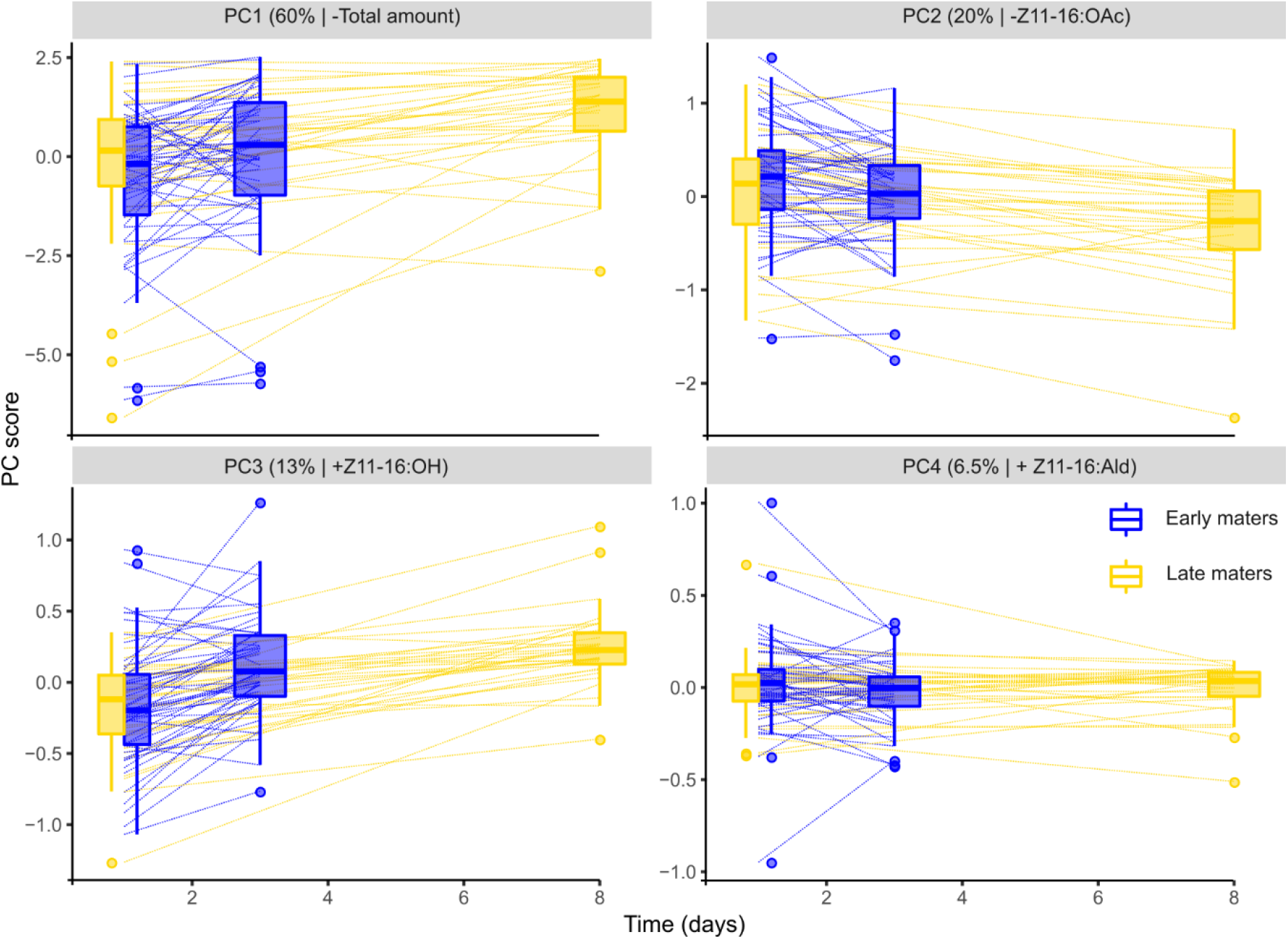
Pheromone variation within and between females. Box-and-whisker plots show distribution of principal component (PC) scores at day one and at day three or eight for early (blue) and late (yellow) maters, respectively. Dashed lines connect samples taken from the same individual. For each PC, the amount variance explained and the main pheromone component that loads on the PC, as well as the sign of the loading (positive, +, or negative, −), is indicated above each panel.

### Relationship between fitness and *inter*-individual pheromone variation

When we determined correlations between the female pheromone signal and her fecundity, fertility and life span, we found that all fitness measurements depended on variation in the pheromone signal. In all models, both in early and late maters, we found at least one PC describing variation in the composition of the pheromone blend to explain variation in fitness (Fig 3A). For fecundity (in both early and late maters) we also observed a correlation with PC1. Partial pseudo-R^2^ values ranged from three to thirty percent for the pheromone components. Time spent calling before mating and the pupal mass of the female also explained fitness variation in most models and the combined effect from the PCs, covariates, and their interactions explained between 15 – 59% of the variation in fitness measurements (Table S2).

**Fig 3.**
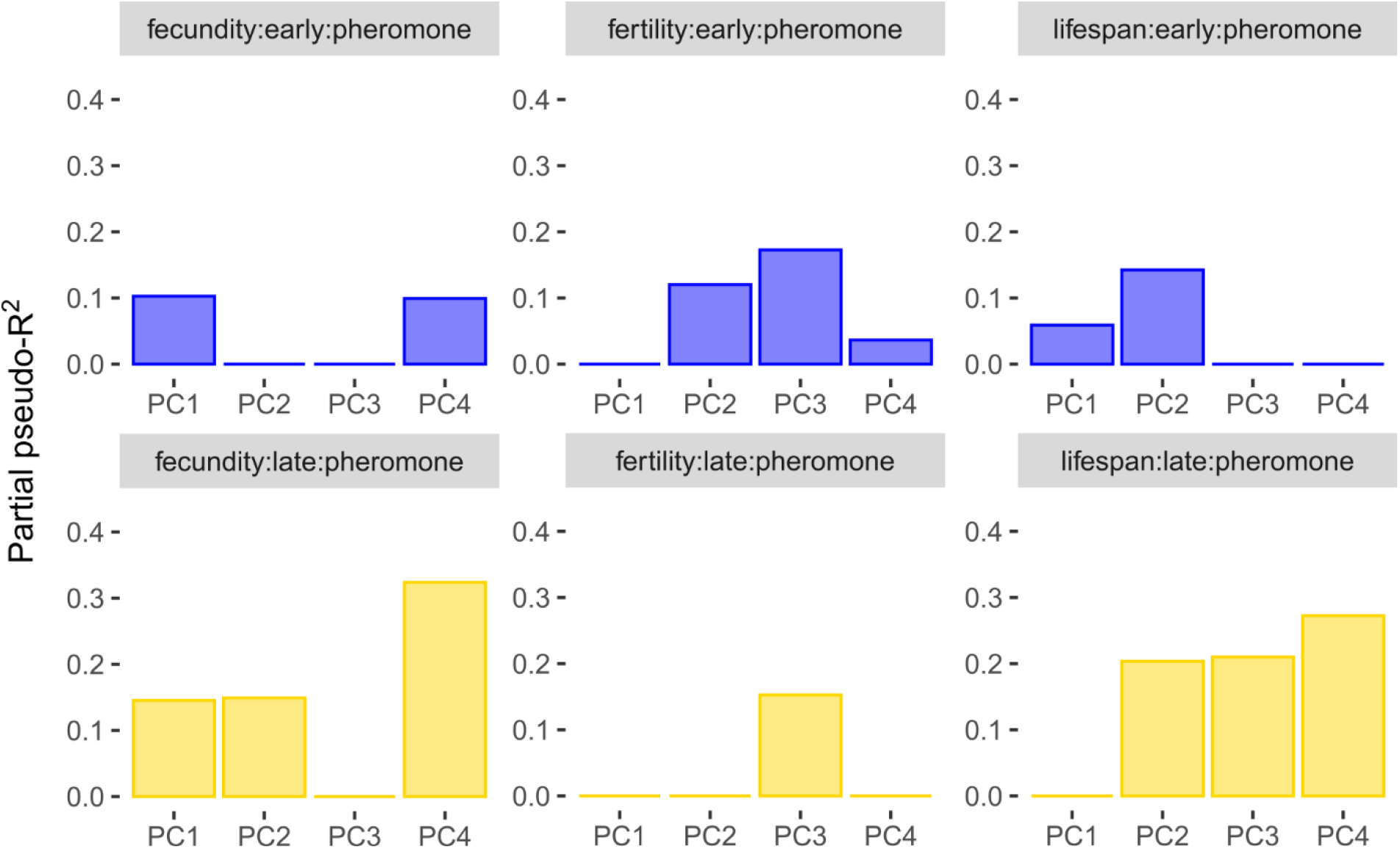
Fitness variation explained by pheromone traits. Each panel depicts one model for either early (blue) or late maters (yellow), for fecundity, fertility, or lifespan as a response variable, and for pheromone amount and composition as a predictor. For each of the four PCs, columns depict the partial pseudo-R^2^ of the PC and all interactions in which the PC is involved in the model.

Producing more pheromone (lower scores on PC1) or higher relative amounts of Z11-16:OAc (lower scores on PC2), Z11-16:OH (higher scores on PC3), and Z11-16:Ald (higher scores on PC4) was universally associated with higher fitness (higher fecundity and fertility, longer lifespan) with one exception: higher scores for PC4 were associated with lower fecundity in the early maters (Fig 4; Fig S4; Table S2). In the case of interactions between a pheromone PC and female pupal mass, we examined the modulating effect of pupal mass on the correlation between fitness and pheromone by drawing separate regression lines for the females with the lowest 33% pupal mass, those with pupal mass in the middle tercile, and those in the top 33% of pupal mass, using the r-package “interactions” (59). This showed that higher relative amount of Z11-16:OH (higher PC3 score) was associated with higher fertility in the heaviest 33% females, but not in the bottom 67% (Fig 4). Also, higher relative amount of Z11-16:OAc (lower PC2 score) was associated with longer life span in heavy females, but with lower life span in lighter females (Fig 4).

**Fig 4.**
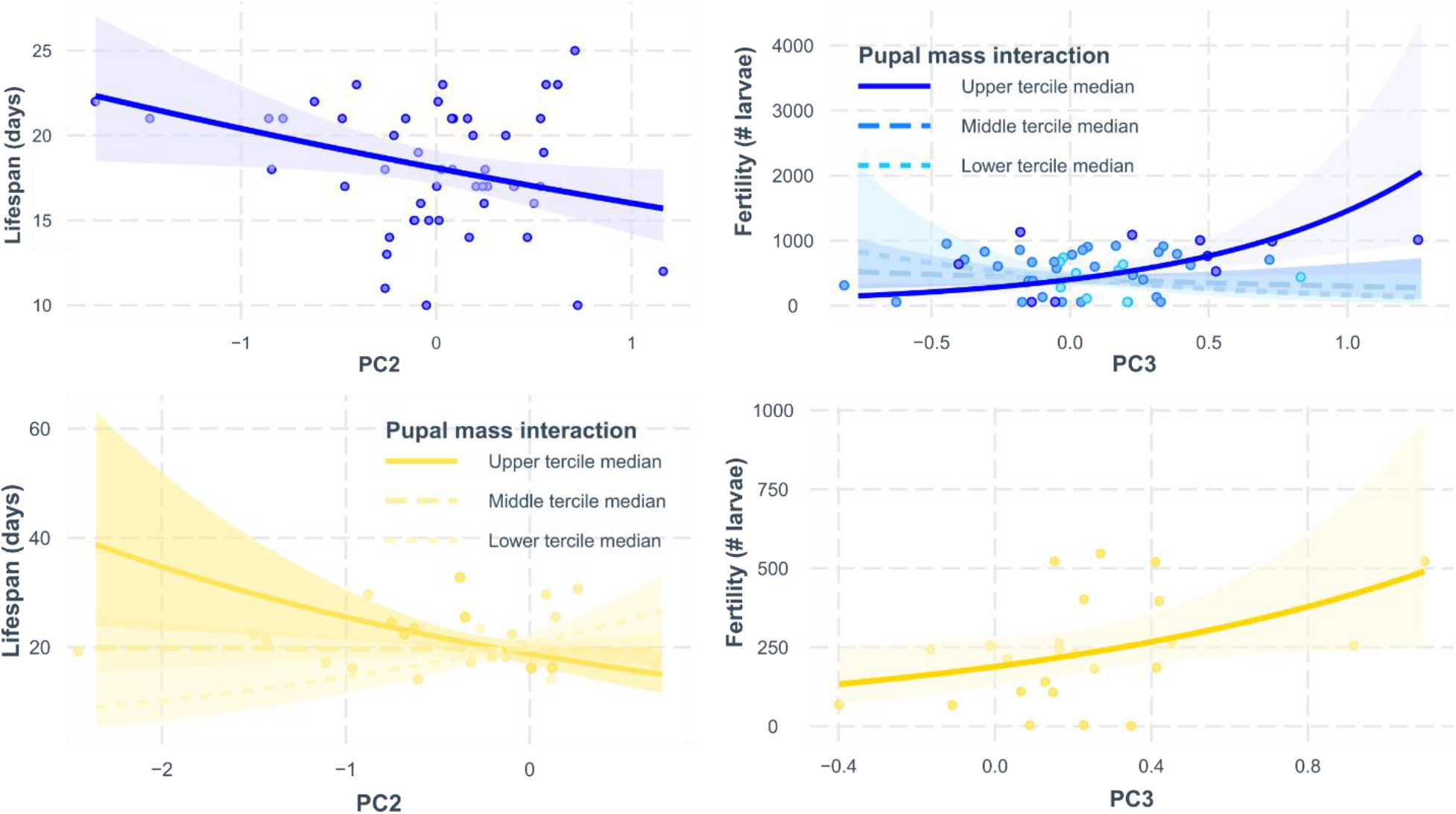
Relationship between pheromone variation and fitness, showing the correlation between PC scores and fitness from both early maters (blue) and late maters (yellow). Points show individual coordinates, lines show regression paths, i.e. the effect of the PC on the x-axis on the response on the y-axis, accounting for the effects from all other predictors in the model. Higher scores on PC2 correspond to lower relative amounts of Z11-16:OAc, while higher scores on PC3 correspond to higher relative amounts of Z11-16:OH. Interaction effects are shown as regression paths split by terciles. For the PCs where we found interaction effects between pupal mass and fitness measures, relationships were determined separately for pupal masses in the lower tercile (median of the lower tercile = 0.224 g), middle tercile (median = 0.238 g), and upper tercile (median = 0.275 g).

### Relationship between fitness and *intra*-individual variation

To test the hypothesis that females with stable pheromone signals during their life time have higher fitness, we calculated intra-individual variation as the absolute difference between PC scores for pheromone measurements taken 24 hours post-eclosion and for pheromone measurements taken three or eight days post-eclosion, for early and late maters respectively. We found that between 6% and 45% of the variation in fitness measures was predicted by intra-individual variation in either the total amount (PC1), the relative amount of Z11-16:OAc (PC2) or both (Fig 5).

**Fig 5.**
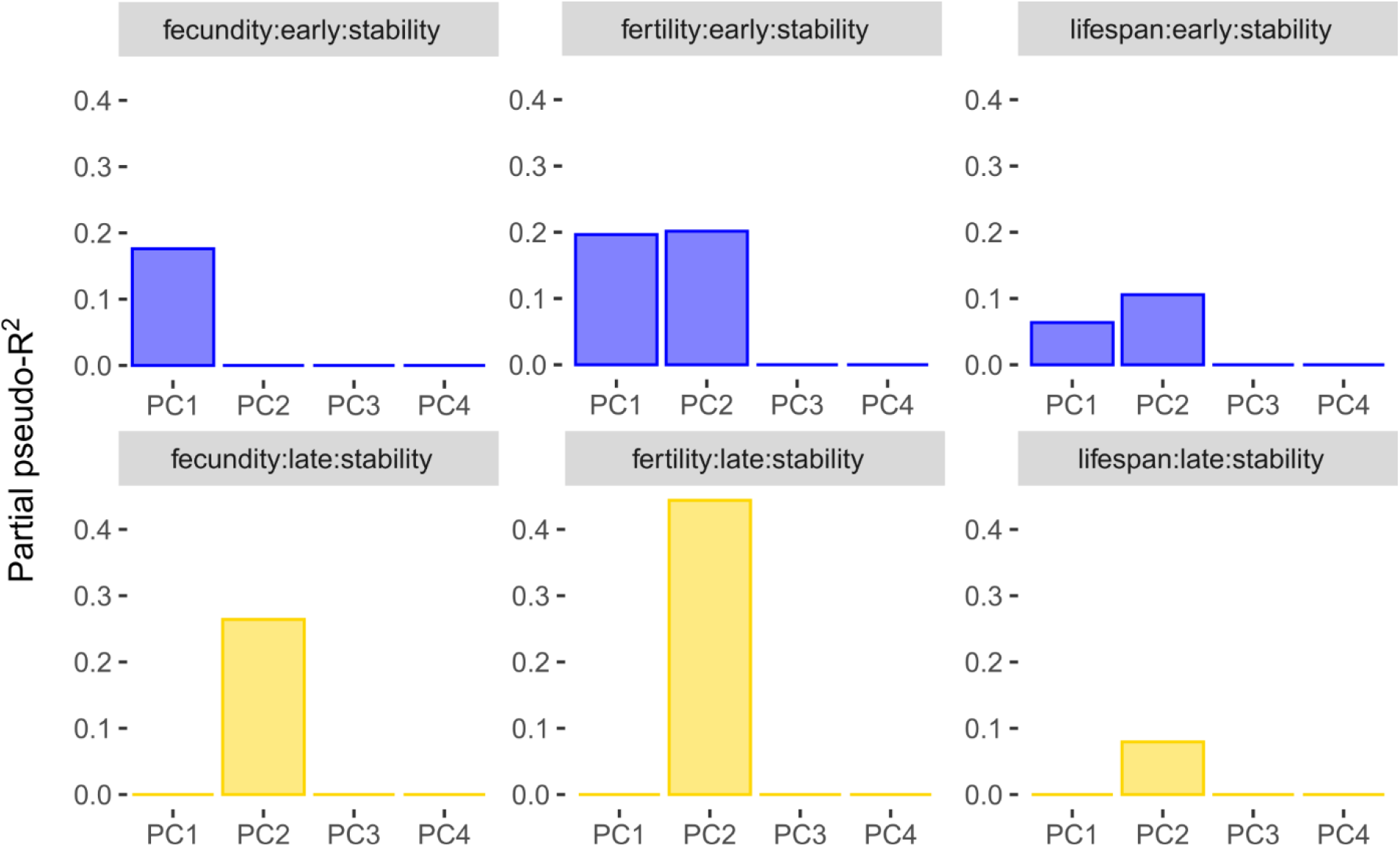
Fitness variation explained by intra-individual variation in pheromone amount and composition (here referred to as stability). Each panel depicts one model for either early (blue) or late maters (yellow), for fecundity, fertility, or lifespan as a response variable, and for intra-individual variation in pheromone PCs as a predictors. Column height corresponds to the partial pseudo-R^2^ of intra-individual variation in the PC and all interactions in which the PC is involved in the model. PCs absent from the model are shown as partial pseudo-R^2^ = 0.

In early maters, more stability in the total amount (PC1) was associated with higher fecundity, fertility, and longevity, while in late maters stability in total amount was uncoupled from fitness (Fig 6; Fig S5). For PC2, we similarly found a predominantly positive correlation between stability and fitness (i.e. lower values on the plasticity axis are associated with higher values on the fitness axis). We also observed examples of effects in the opposite direction for both fertility and life span (Fig 6; Fig S5).

**Fig 6.**
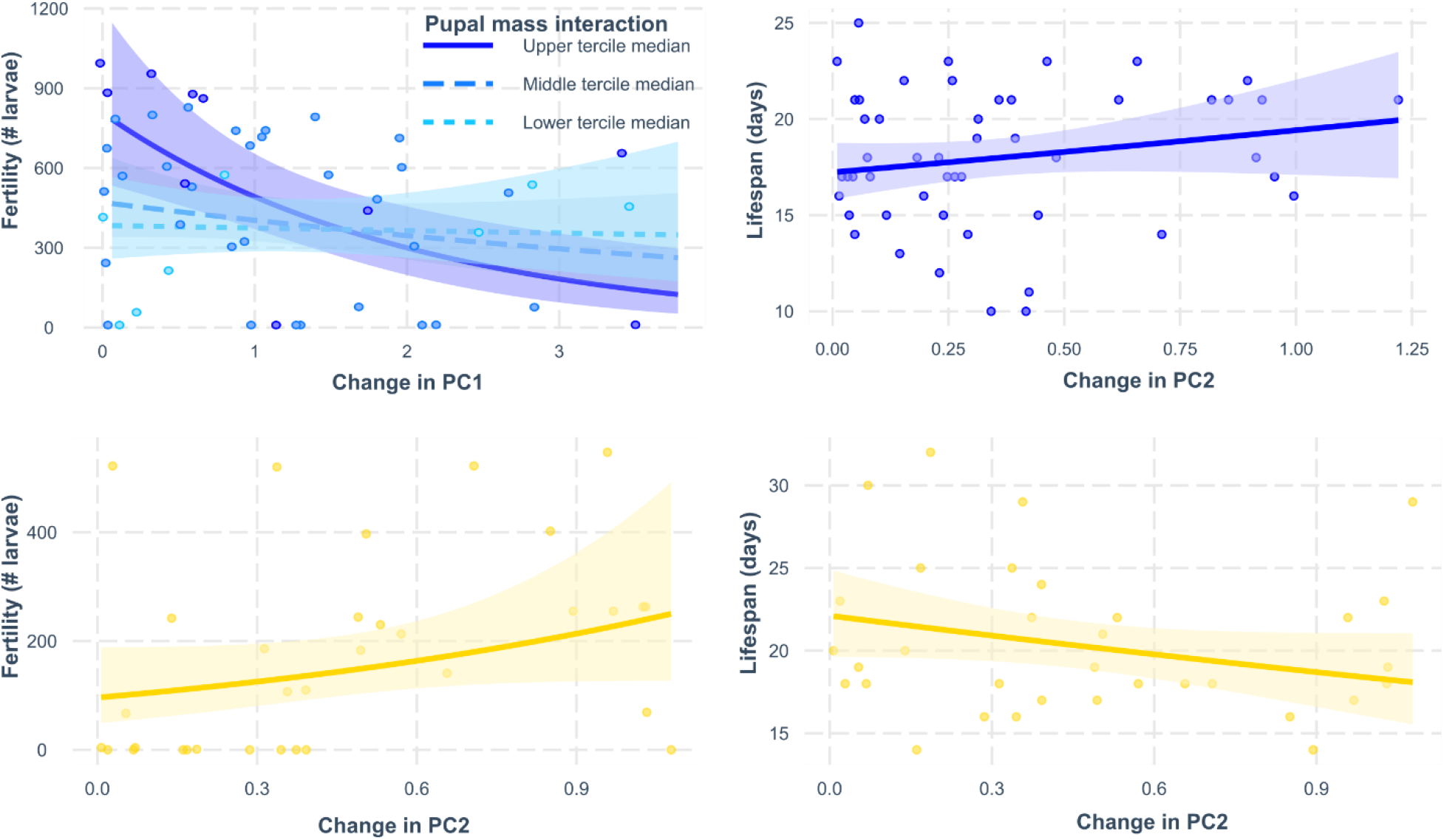
Relationship between change in pheromone and fitness, showing the magnitude and direction of the correlation between change in PC scores and fitness in early maters (blue) and late maters (yellow). Change in PC scores was calculated as the absolute difference between the PC score for the first sample and second sample of an individual. Points show individual coordinates, lines show regression paths, i.e. the effect of the PC on the x-axis on the response on the y-axis, accounting for the effects from all other predictors in the model. For pupal mass, the lower tercile median = 0.224 g, the middle tercile median = 0.238 g, the upper tercile median = 0.275 g.

## DISCUSSION

If sex pheromone signals are costly, calling activity, pheromone amount and/or pheromone composition should covary with fitness. We showed that in *H. subflexa* an extreme delay in mating of 8 days and a continued investment in signaling was associated with lower reproductive output. We also found that while virgin females maintained high signaling activity, their signal changed over time: the total amount decreased and the ratios of components changed. Signal variation was proportional to the delay in mating, but varied considerably across females. Some females showed high intra-individual variation, while others had stable signals over time. The heritability estimates of the intra-individual variation in pheromone composition (roughly between 0.1 and 0.2) indicated that this variation can evolve in response to selection. Lastly, we found that longevity, fecundity, and fertility were correlated with pheromone amount, with pheromone composition, and with the stability in both pheromone amount and composition within females. We thus conclude that our results meet expectations for costly pheromone signaling and that the amount, composition, and maintenance of the long-distance mate attraction pheromone produced by *H. subflexa* females likely depend on the genetic quality of the female.

### Calling activity and fitness

In a synthesis of empirical results on calling activity in female moths, the majority of moth species were found to increase their calling effort over time (49). This finding is in line with theoretical predictions for a costly signal, for which the production depends on the number of male arrivals: a virgin female should not invest much in signaling if this draws in too many males, but she should increase signaling efforts if no males visit at all (49). Our results show that for *H. subflexa,* calling activity initially indeed increased, but then decreased (Fig 1). Females that spent more nights calling had reduced reproductive success, but slightly longer life span relative to females that called fewer days, which is in line with general patterns across moths and suggest both time constraints (fewer days remaining to lay eggs in late mated females) and energy constraints (e.g., resorption of eggs to support somatic maintenance) to sex pheromone calling (60). Together, these findings support our hypothesis that there are costs associated with sex pheromone calling in female moths.

### Pheromone variation and fitness

We hypothesized that these costs would manifest in signal-fitness correlations in virgin moths. We found evidence for covariation between fitness and sex pheromone amount, composition, and stability, regardless of the fitness measure (reproductive output or life span). In addition, we found similar patterns in two independent groups of females that differed in their age at the time of sampling and in their remaining life span to lay eggs. Even though in nature females are unlikely to stay virgin for 8 days (late maters), our finding that her signal still correlates with fitness shows that the patterns that we found are independent of female age. Our findings thus strongly support a scenario for costly sex pheromone signals in moths and provide context to earlier findings that suggested condition-dependence of moth sex pheromone amount (38,39) and composition (40), by revealing positive covariance between signal and fitness components.

On the one hand, the consistent positive correlations between fitness and pheromone amount as well as between fitness and the relative abundance of pheromone components important to male mate attraction are surprising from a physiological perspective. It is unlikely that pheromone production presents significant metabolic costs to signaling females, because the nutrients used for pheromone production are negligible compared to available resources (27). However, there may be indirect costs, for example if enzymes that convert sex pheromone component precursors into their final products (in the appropriate ratios of species-specific blends) are also used for other critical physiological processes.

On the other hand, the fitness costs to sex pheromone composition inferred here can explain field observations of sex pheromone variation among wild populations of *H. subflexa*. The acetate ester Z11-16:OAc slightly improves the attractiveness of the *H. subflexa* blend, while strongly deterring the closely-related tobacco budworm, *H. virescens* (61,62). In regions where *H. virescens* co-occurs with *H. subflexa*, the relative amount of Z11-16:OAc of field-caught females is significantly higher compared to regions where *H. virescens* is absent (63). Apparently, Z11-16:OAc is only produced when communication interference is likely to happen. As we found that the PC2 axis, associated with variation in Z11-16:OAc, explained 10-20% of the variation in fecundity, fertility, and life span (Fig 3), such that females with higher fitness had higher relative amounts of Z11-16:OAc, selection likely favors females that produce lower relative amounts of acetate esters when there is not risk of heterospecific mate attraction.

There is limited data available for male response to intra-specific levels of variation in moth sex pheromones and these data often come from different methods, including sticky traps, field traps and wind tunnel assays. It is therefore difficult to judge the relevance of the levels of observed variation to male response. The two axes that described most of the variation in pheromone composition (and that correlated with fitness), PC2 and PC3, are driven by variation in Z11-16:OAc, Z11-16:OH, and Z9-16:Ald, which all have been shown to be important for male attraction (61,62). The relative amount of Z11-16:OAc ranged from 5% to 15%. Previous studies have shown that a) as little as 1% is enough to completely deter the heterospecific *H. virescens* (64), b) adding Z11-16:OAc to a synthetic blend increases the attractivity of *H. subflexa* males (61,65), and c) increased levels of acetate esters (from roughly 1-3 % to 10-15%) also increased male response (62). For Z11-16:OH, the lowest and highest scores on PC3 corresponded to a 2-fold difference, i.e. from ~3% to ~6%. Previous studies have shown that increasing or reducing the relative amount of Z11-16:OH by a factor 2 can reduce male attraction to sticky traps by 10% in the field (66), while in wind tunnel experiments a change in male attraction was not found (61). Finally, for Z9-16:Ald we found a range from ~15% to ~35%. Previous studies have found a 38% reduction in male precopulatory behavior when relative amounts of Z9-16:Ald were below 15% (67), while in wind tunnel assays Z9-16:Ald levels below 10% reduced male attraction by 20% (61). Hence, the observed levels of variation in the pheromone summarized by PC2 and PC3 fall within a range that is likely relevant for variation in male responses.

We also hypothesized that for costly signals, maintenance is costly too and we thus expected stability of the signal to covary with fitness. We found that individuals with more stable sex pheromone signals over time had higher fitness, although negative correlations were found as well. The evidence thus supports our hypothesis for fitness costs to sex pheromone stability. Since in moths older females are typically less attractive, as measured by mating success (42–48), it is possible that a female benefits from maintaining a young-female-like blend. This would be similar to cricket song and mouse urinary protein pheromones, for which signal composition reliably reflects age and in both cases the scent of senescence is associated with reduced mate attraction (68,69). Additionally, female moths produce more pheromone earlier in life. Possible physiological explanations for decreasing pheromone amounts in aging females are that changes in Juvenile Hormone may result in hormonal suppression of pheromone production later in life (70,71) or that older females may have a reduced capability to synthesize pheromone components (71). Lower pheromone amounts can reduce the attractiveness of the blend, however there is mixed evidence for this hypothesis (49). Therefore, an evolutionary mechanism that would explain a benefit to costly stability is that stability counters signal decay due to senescence.

In summary, we find evidence for signal-fitness covariation, which thus indicate costs to sex pheromone signaling. Our results are in line with earlier findings for condition-dependence of pheromone amount in moths and beetles, but go beyond these findings by revealing (i) a relationship between a moth sex pheromone signal and fitness and (ii) finding this relationship not only for the amount of pheromone produced, but also for the composition and for the extent to which females keep their signal amount and composition stable. These results add multiple new dimensions in which genetic quality differences between female moths can contribute to pheromone variation within species. As genetic variation underlying sexual signals ultimately provides the raw material from which species barriers may originate, the relationship between signal variation and fitness components presented here thus provides a mechanism for the evolution of sex pheromone signals and the diversity of moth species.

## ACKNOWLEDGEMENTS

The authors thank the Evolutionary and Population Biology group members, especially Isabel Smallegange and Emily Burdfield-Steel for their helpful discussions and the editorial board member and three referees for their constructive criticism. This project is funded by the Netherlands Organization for Scientific Research (NWO-ALW, award no. 822.01.012 awarded to ATG), the National Science Foundation (award no. IOS-1052238 and IOS-1456973 awarded to MW), and by the Marie Skłodowska-Curie Individual fellowship (grant agreement No 794254 awarded to TB).

