## Supplementary Figures for "Sex pheromone signal and stability covary with fitness"

### Fig S1 Time of onset of calling

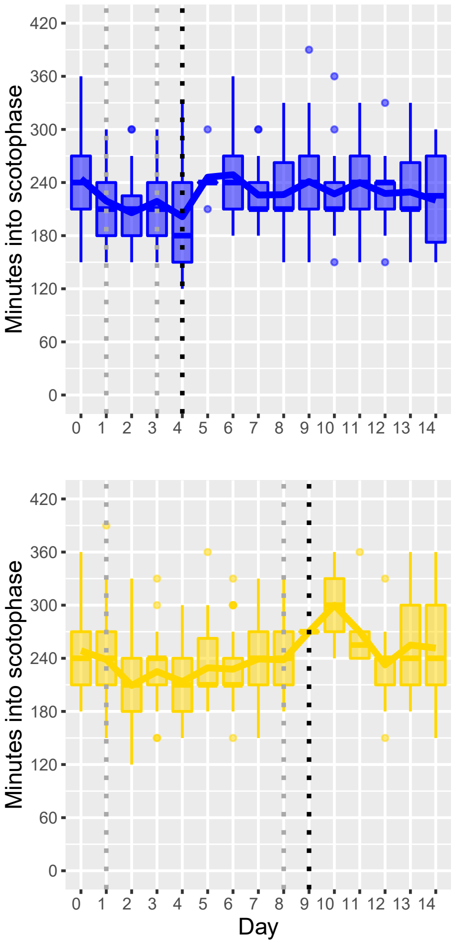

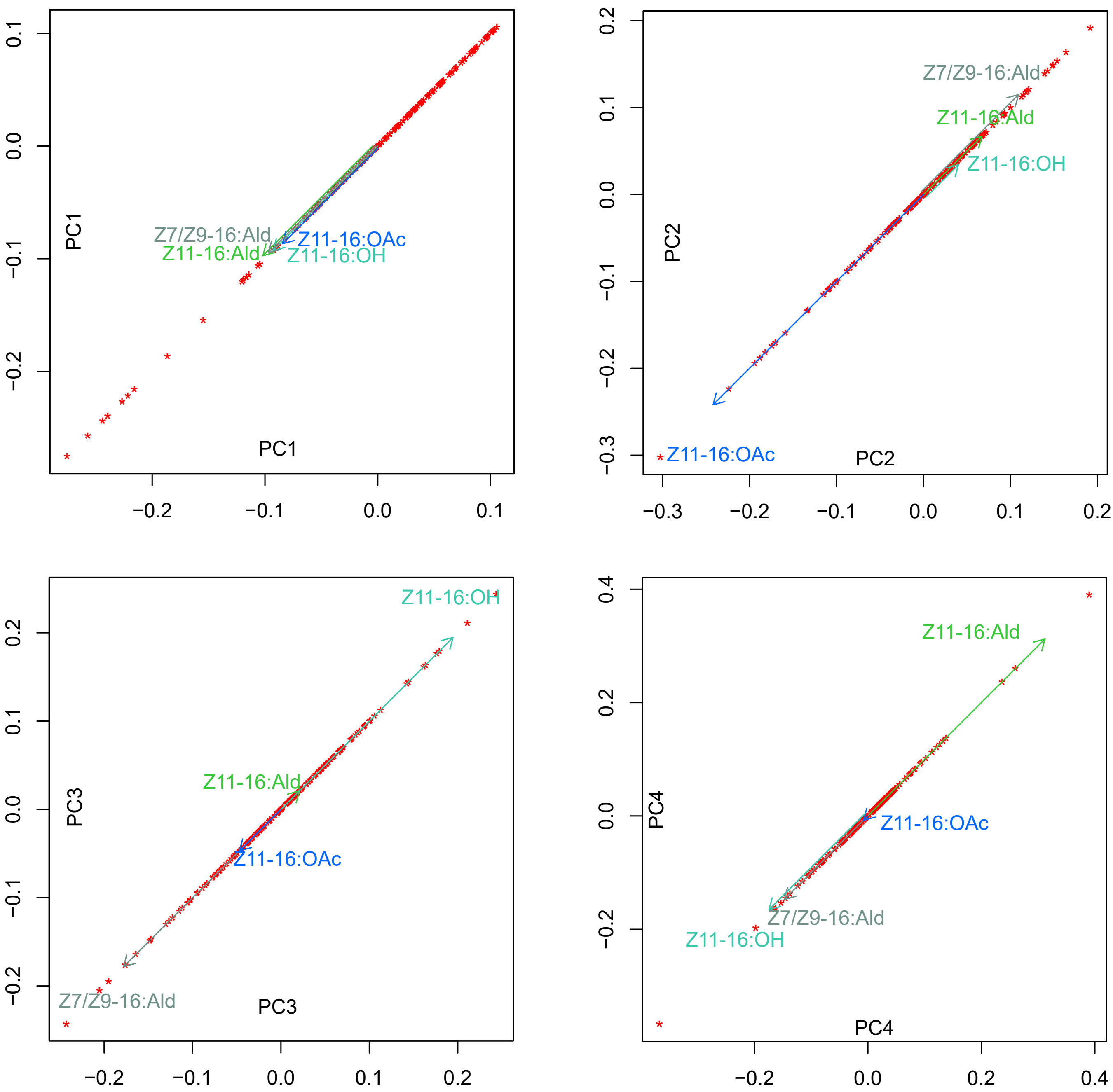

PC1
Z7/Z9-16:Ald
Z11-16:Ald
Z11-16:OAc
Z11-16:OH
PC2
Z7/Z9-16:Ald
Z11-16:Ald
Z11-16:OH
Z11-16:OAc
PC3
Z11-16:OH
Z11-16:Ald
Z11-16:OAc
Z7/Z9-16:Ald
PC4
Z11-16:Ald
Z11-16:OAc
Z7/Z9-16:Ald
Z11-16:OH

Narrow-sense heritability calculated using MCMCglmm with inverse gamma prior

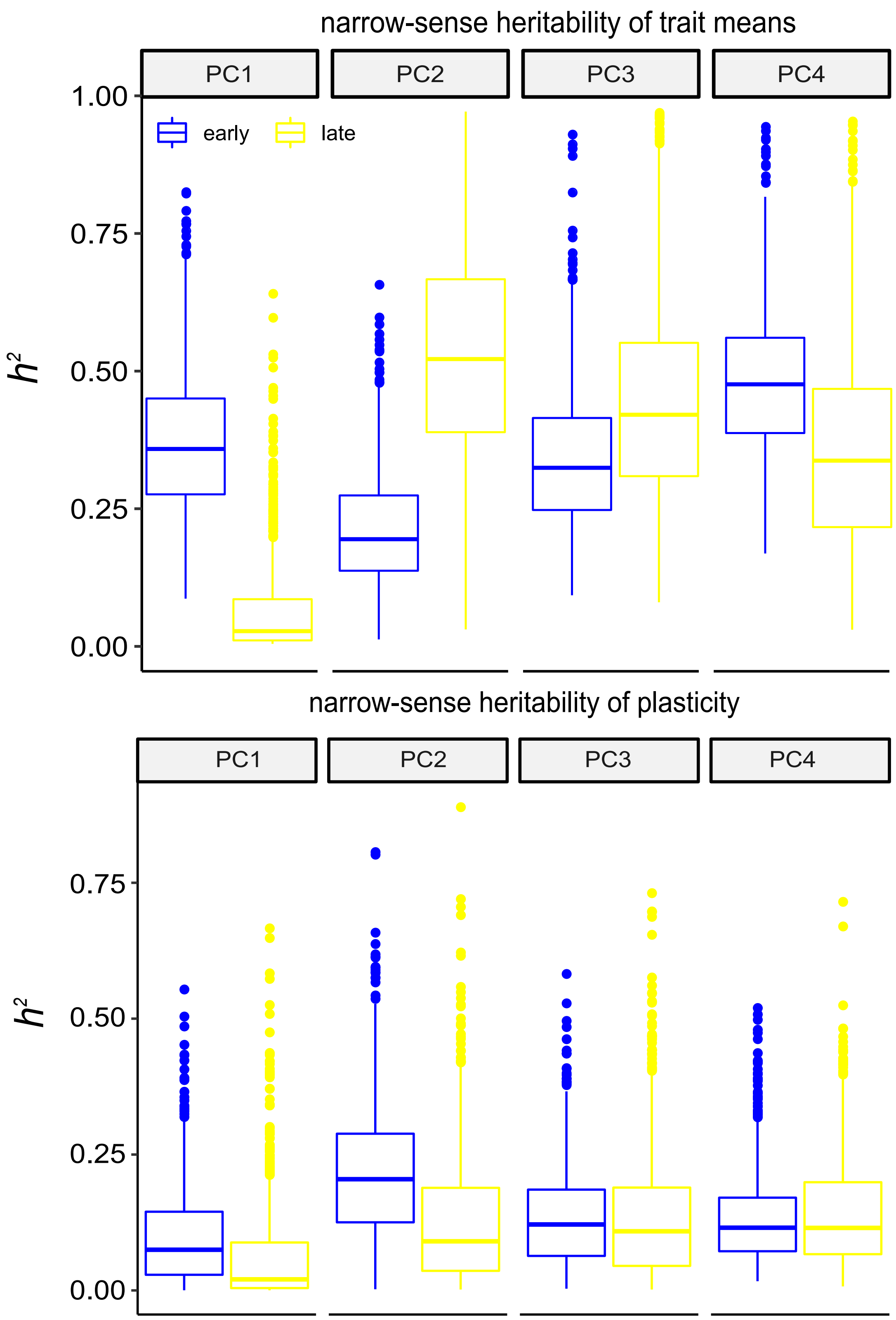

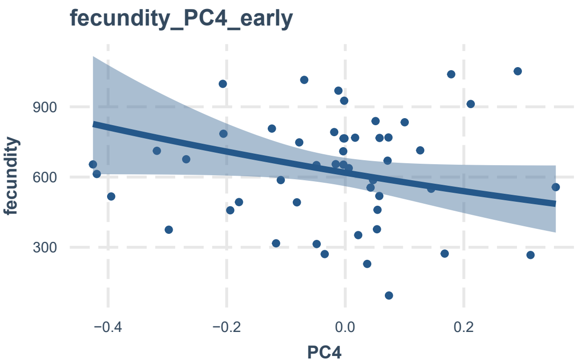
fecundity_PC4_early
fecundity
900
600
300
−0.4
−0.2
0.0
0.2
PC4

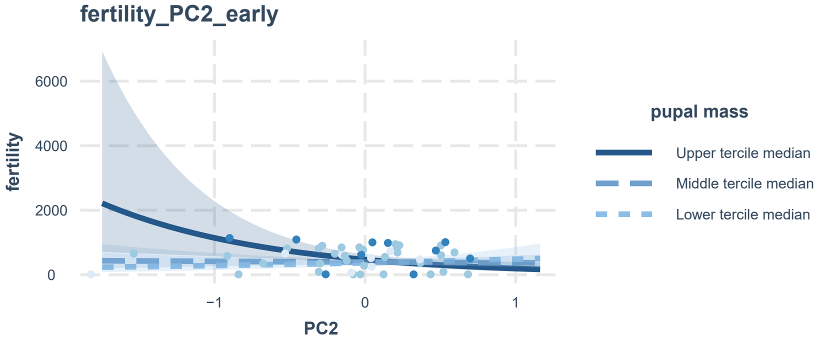
fertility_PC2_early
fertility
6000
4000
2000
0
−1
0
1
PC2
pupal mass
Upper tercile median
Middle tercile median
Lower tercile median

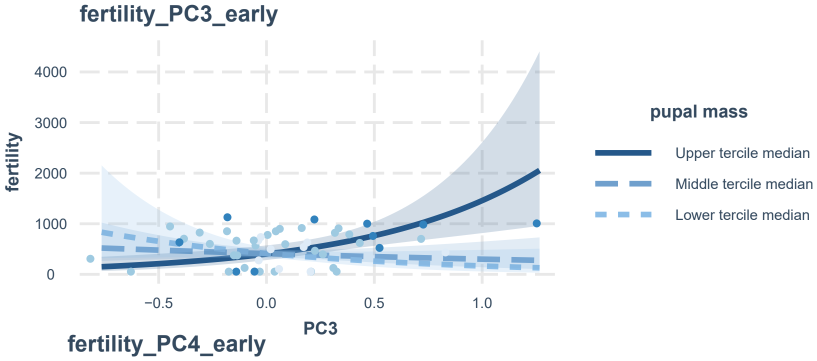
fertility_PC3_early
fertility
4000
3000
2000
1000
0
−0.5
0.0
0.5
1.0
PC3
pupal mass
Upper tercile median
Middle tercile median
Lower tercile median

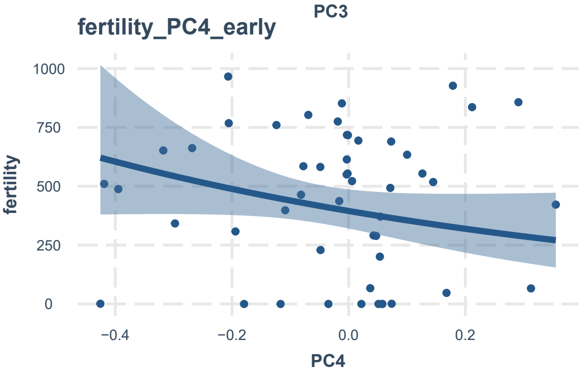
fertility_PC4_early
fertility
1000
750
500
250
0
−0.4
−0.2
0.0
0.2
PC4

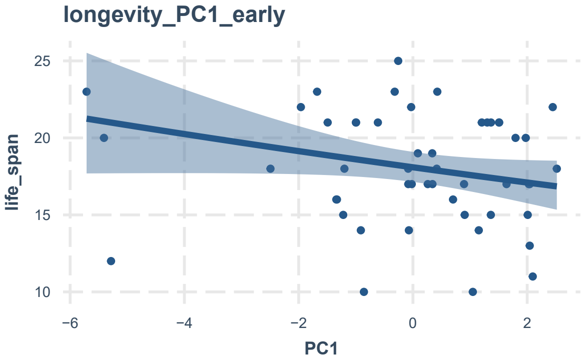
longevity_PC1_early
life_span
25
20
15
10
−6
−4
−2
0
2
PC1

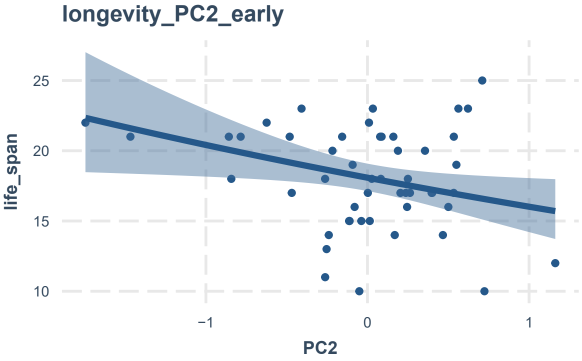
longevity_PC2_early
life_span
25
20
15
10
−1
0
1
PC2

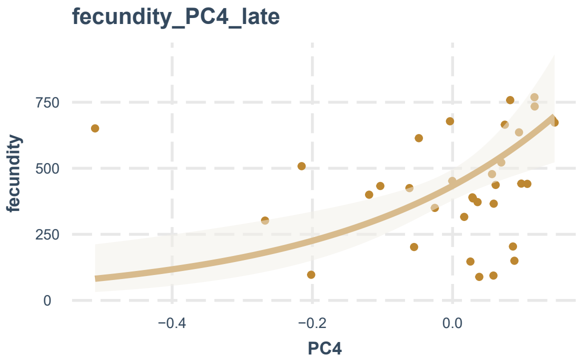
fecundity_PC4_late
fecundity
PC4

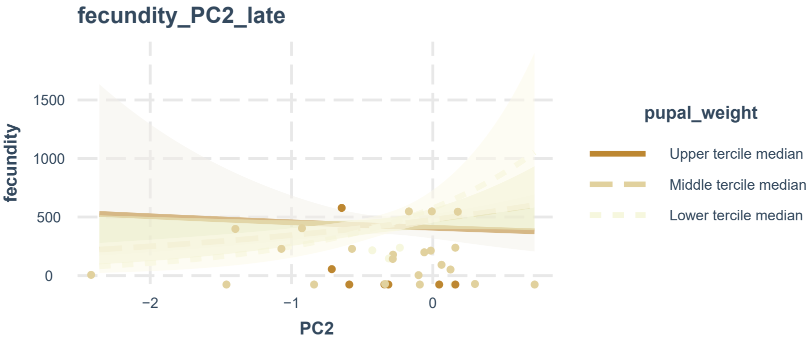
fecundity_PC2_late
fecundity
PC2
pupal_weight
Upper tercile median
Middle tercile median
Lower tercile median

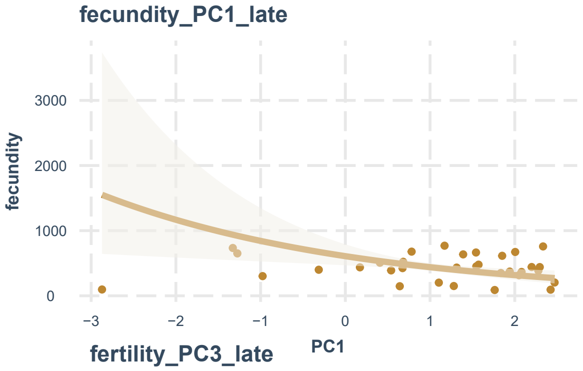
fecundity_PC1_late
fecundity
PC1

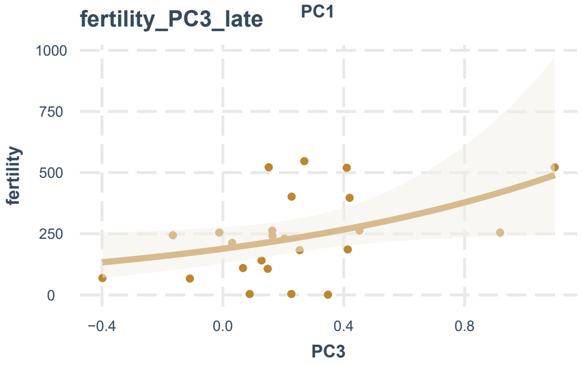
fertility_PC3_late
fertility
PC3

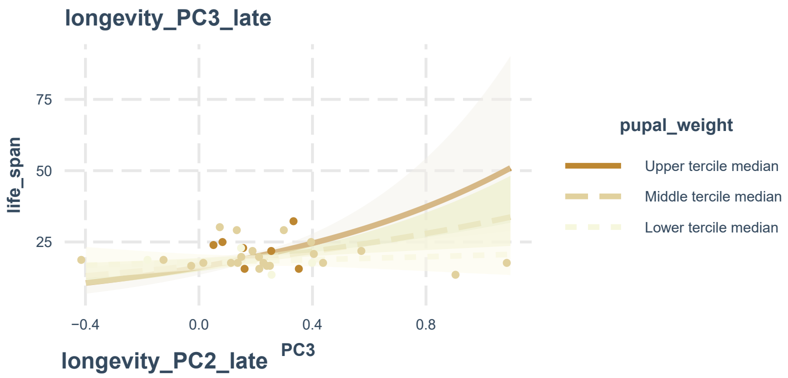
longevity_PC3_late
life_span
PC3
pupal_weight
Upper tercile median
Middle tercile median
Lower tercile median

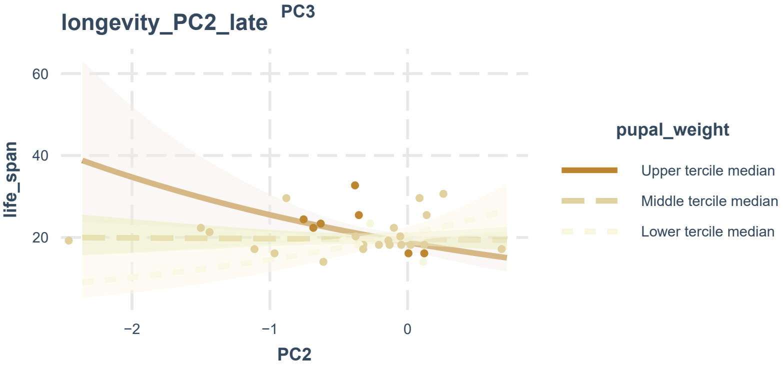
longevity_PC2_late
life_span
PC2
pupal_weight
Upper tercile median
Middle tercile median
Lower tercile median

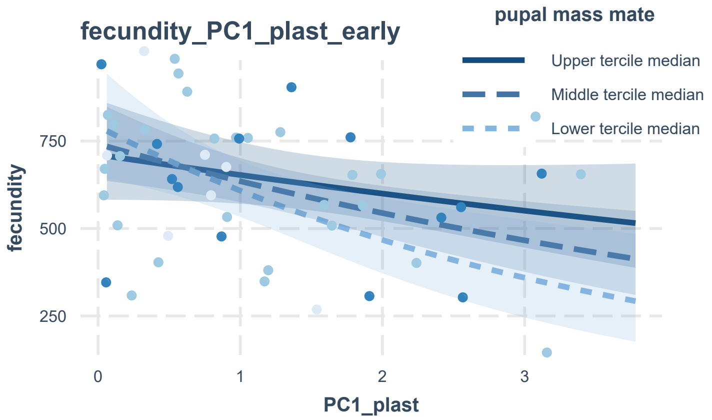
fecundity_PC1_plast_early
pupal mass mate
Upper tercile median
Middle tercile median
Lower tercile median
fecundity
750
500
250
0
1
2
3
PC1_plast

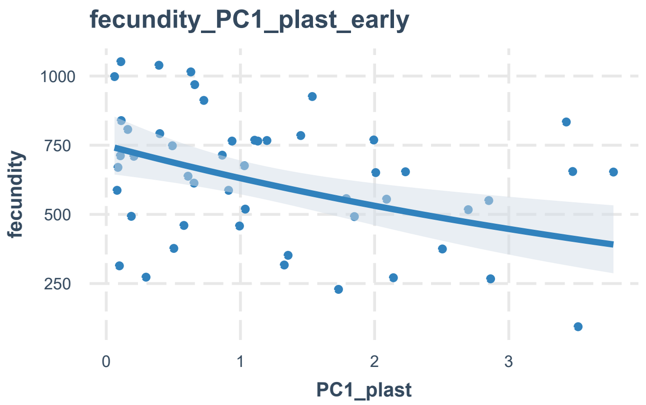
fecundity_PC1_plast_early
fecundity
1000
750
500
250
0
1
2
3
PC1_plast

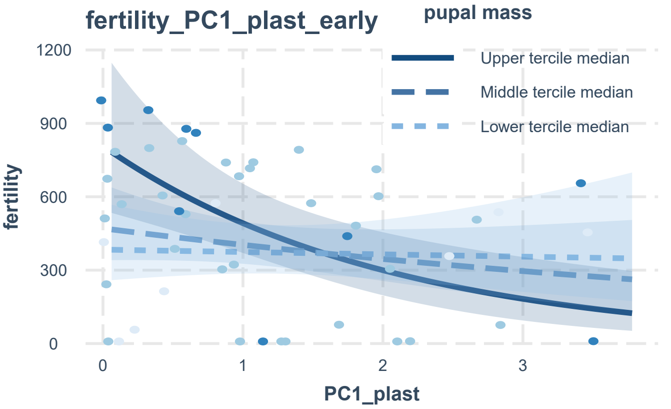
fertility_PC1_plast_early
pupal mass
Upper tercile median
Middle tercile median
Lower tercile median
fertility
1200
900
600
300
0
0
1
2
3
PC1_plast

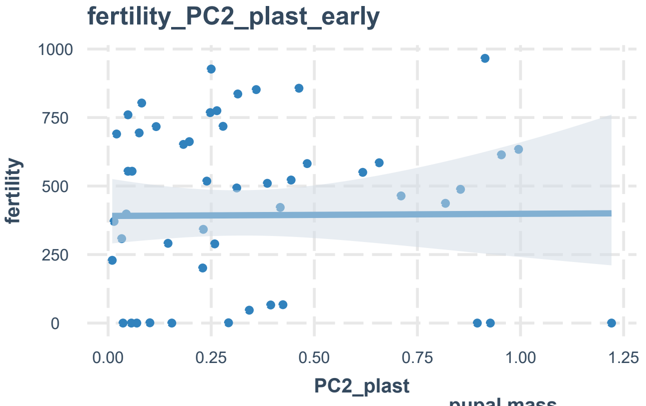
fertility_PC2_plast_early
fertility
1000
750
500
250
0
0.00
0.25
0.50
0.75
1.00
1.25
PC2_plast

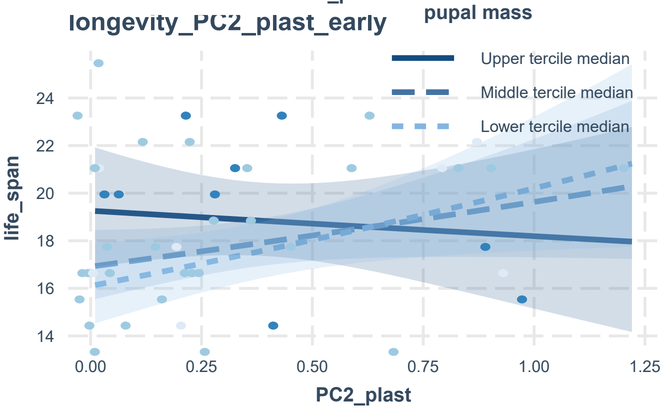
longevity_PC2_plast_early
pupal mass
Upper tercile median
Middle tercile median
Lower tercile median
life_span
24
22
20
18
16
14
0.00
0.25
0.50
0.75
1.00
1.25
PC2_plast

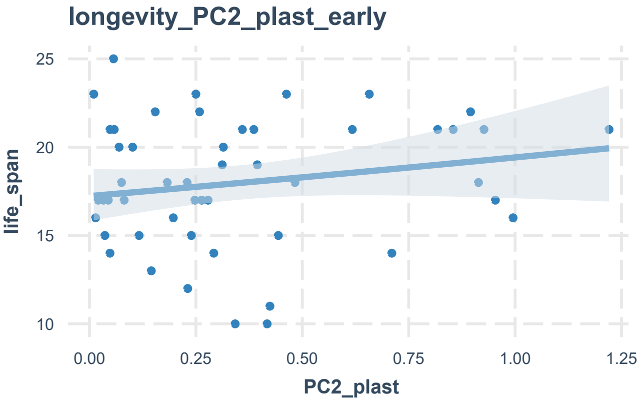
longevity_PC2_plast_early
life_span
25
20
15
10
0.00
0.25
0.50
0.75
1.00
1.25
PC2_plast

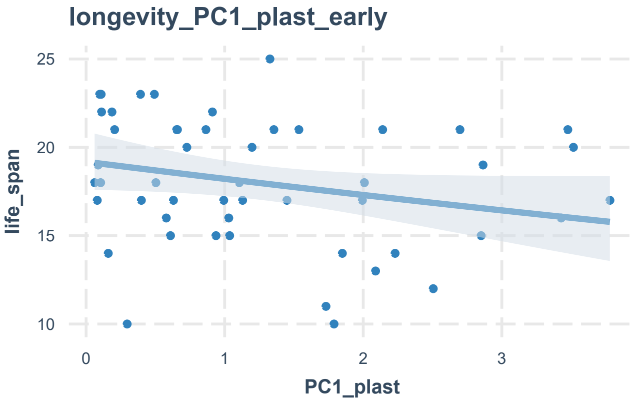
longevity_PC1_plast_early
life_span
25
20
15
10
0
1
2
3
PC1_plast

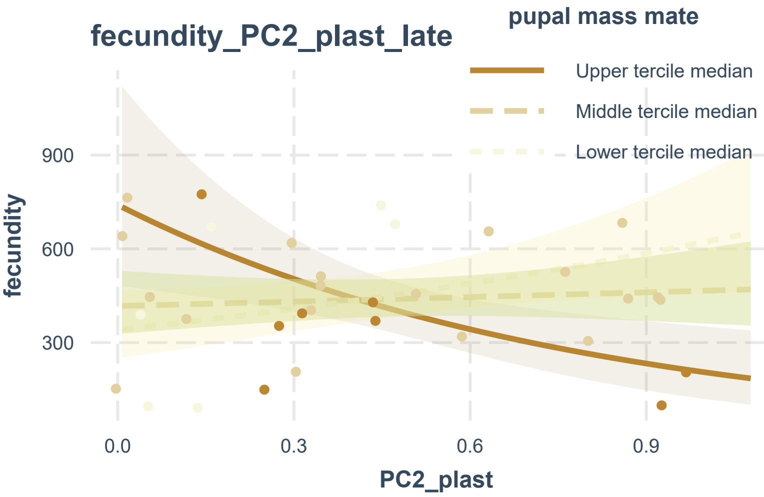
fecundity_PC2_plast_late
pupal mass mate
Upper tercile median
Middle tercile median
Lower tercile median
fecundity
900
600
300
0.0
0.3
0.6
0.9
PC2_plast

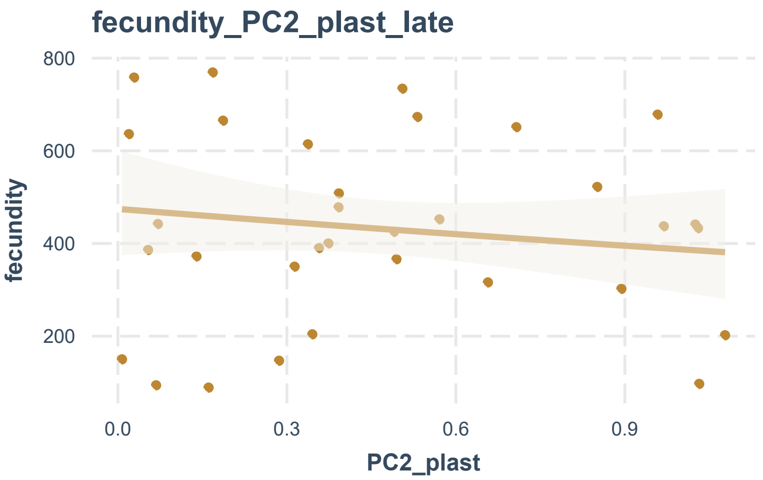
fecundity_PC2_plast_late
fecundity
800
600
400
200
0.0
0.3
0.6
0.9
PC2_plast

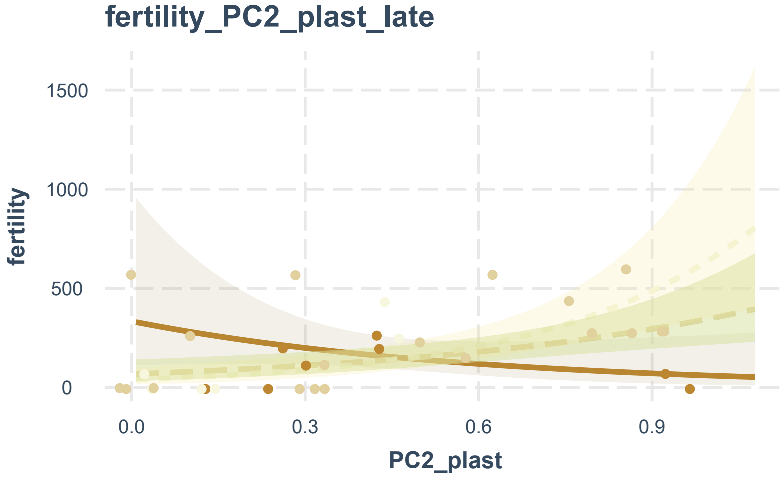
fertility_PC2_plast_late
fertility
1500
1000
500
0
0.0
0.3
0.6
0.9
PC2_plast

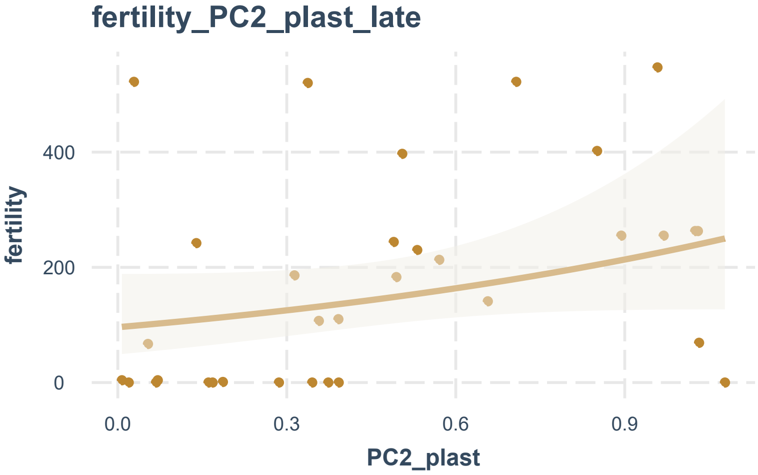
fertility_PC2_plast_late
fertility
400
200
0
0.0
0.3
0.6
0.9
PC2_plast

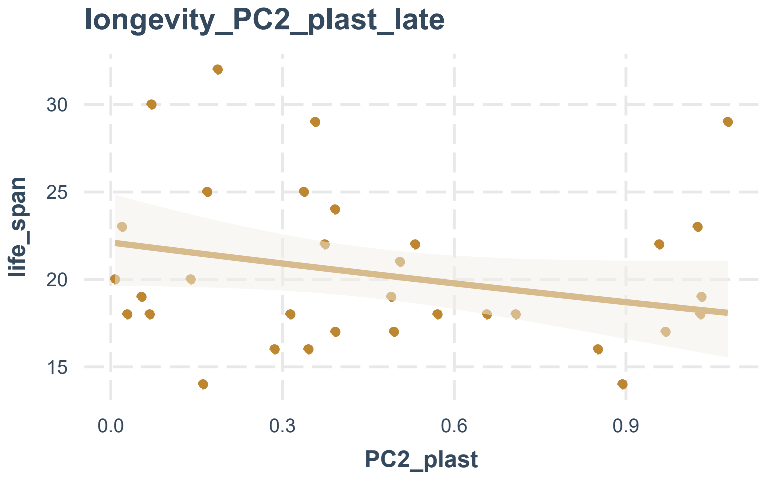
longevity_PC2_plast_late
life_span
30
25
20
15
0.0
0.3
0.6
0.9
PC2_plast
